# Surface Generative Modelling of Neurodevelopmental Trajectories

**DOI:** 10.1101/2023.10.16.562598

**Authors:** Abdulah Fawaz, Saga N. B. Masui, Logan Z. J. Williams, Simon Dahan, A. David Edwards, Emma C. Robinson

**Affiliations:** Department of Biomedical Engineering, School of Biomedical Engineering and Imaging Sciences, King’s College London, UK; Centre for the Developing Brain, School of Biomedical Engineering and Imaging Sciences, King’s College London, UK; Department for Forensic and Neurodevelopmental Sciences, and the MRC Centre for Neurodevelopmental Disorders, King’s College London, UK

**Keywords:** Graph Neural Network, GAN, cortical surface, generative modelling, neurodevelopment

## Abstract

Cortical neurodevelopment is sensitive to disruption following preterm birth, with lasting impact on cognitive outcomes. The creation of generative models of neurodevelopment could aid clinicians in identifying at-risk subjects but is complicated by the degree of subject variability in cortical folding, and significant heterogeneity in the effect of preterm birth. In this work, we propose a graph convolutional generative adversarial network (GAN) and a training scheme to simulate neonatal cortical surface developmental trajectories. The proposed model is used to smoothly modify two cortical phenotypes: post-menstrual age at scan (PMA) and gestational age at birth (GA) on data from the developing Human Connectome Project (dHCP) [1]. The synthetic images were validated with an independently trained regression network, and compared against follow up scans, indicating that the model can realistically age individuals whilst preserving subject-specific cortical morphology. Deviation between simulated ‘healthy’ scans, and preterm follow up scans generated a metric of individual atypicality, which improved prediction of 18-month cognitive outcome over GA alone.

## I. Background

During the third trimester of pregnancy, the human brain develops significantly in terms of microstructure and functional organisation. This process is known to be sensitive to factors such as preterm birth. Multiple studies have reported significant changes in both cortical microstructure and morphology, including changes to mean diffusivity [2]–[4], cortical thickness and surface area [3], [5]–[7], and reduced white and grey matter volumes [7]–[9]. Atypical cortical neurodevelopment is known to be correlated to poorer cognitive outcomes including Attention-Deficit Hyperactivity Disorder (ADHD), Autism Spectrum Disorders (ASD) and cerebral palsy [10]–[13].

Generative modelling of cortical neurodevelopment could aid clinicians by improving mechanistic understanding of the causes of disease, or be used to identify patients for early clinical intervention. Computational modelling of cortical neurodevelopment faces numerous challenges, particularly due to heterogeneity of cortical organisation and folding patterns, among even healthy subjects. This limits the accuracy of population-based analyses based on diffeomorphic registration, which is insufficient to normalise all sources of variability, and so under which residual misalignments remain [14], [15]. Furthermore, the impact of preterm birth has itself been shown to be heterogeneous [3] - meaning that neurodevelopmental trajectories are unique to individuals.

One increasingly popular approach for modelling heterogeneous brain phenotypes is normative modelling, which attempts to predict typical patterns of neurodevelopment which then act as a reference to understand deviations from the norm. Gaussian Process Regression normative models have found some success in characterising the impact of prematurity on cortical neurodevelopment [16], [17], but remain reliant on image registration leading to high uncertainty at the cortex where intersubject variability is highest [3].

Deep generative modelling has emerged as a powerful tool in overcoming these challenges, leveraging the ability of convolutional neural networks (CNNs) to model complex tasks independent of image registration. Deep generative modelling has already been used to simulate age-related disease progression on volumetric neuroimaging data [18]. For example, Bass, Silva, Sudre, *et al*. [19], [20] used a VAE-GAN [21], with separate content and attribute encodings, to disentangle class specific features of disease, from class irrelevant features of cortical shape variation, to map patterns of disease related brain atrophy in individuals with mild cognitive impairment (MCI) or Alzheimer’s disease (AD). Others models have sought to improve interpretability by incorporating confounding factors, such as age: for example, Ravi, Alexander, Oxtoby, *et al*. [22] proposed a Degenerative Adversarial NeuroImage Net (DaniNet) to simulate AD progression, conditioned on age, disease state and a biological model of disease progression. More recent developments include the use of latent diffusion modelling, to generate age-conditioned brain images with significantly higher perceptual quality than those produced with GANs [23]. Other approaches have sought to go beyond interpretation towards explanation through incorporation of concepts from causal inference [24].

The closest related work to this paper is the age progression/regression model described in Xia, Chartsias, Tsaftaris, *et al*. [25], which uses a GAN, directly conditioned on age difference, to generate subject-specific brain age progression maps (on 2D MRI slices) without the need for longitudinal data. In followup work, the same authors extended their model by further conditioning on health state, to simulate brain ageing with and without MCI or AD [26].

The above models, whilst powerful, are suitable only for 2D images and/or 3D volumes of the brain, yet surface-based analysis of the cortex has been shown to be advantageous over volume-based due to its ability to more accurately encode geodesic distances along the cortex, and hence model patterns of cortical organisation [14], [27]. However, the application of deep learning to surfaces is non-trivial, as surfaces belong to a class of data domains that includes graphs, point clouds and manifolds, which are mathematically inconsistent with foundational aspects of traditional CNNs [28].

Geometric deep learning (gDL) is a nascent field generalising deep learning to a range of non-Euclidean domains. A thorough review of these methods and the theory behind them can be found in the literature [29], but they have already begun to find application on the cortical surface; for cortical surface segmentation [30], [31], regression of phenotypes [32], [33], and cortical surface registration [34], [35]. Comparative studies of gDL models on the cortical surface [33], [36] have highlighted that these models often compromise on either computational feasibility, network expressivity, or fidelity to the symmetry of the underlying data [36]. Generative gDL models have been proposed for drug discovery [37], 3D shape manipulation [38] and point cloud generation [39], but to our knowledge are yet to be applied to generative modelling of cortical imaging features.

In this paper we propose an extension to our preliminary work [40], which trained a CycleGAN to translate cortical appearance of preterm scans to appear like healthy term controls. As the model was conditioned purely on discrete classes, it was was not able to *continuously* age brains. This work therefore proposes a graph convolutional GAN, capable of simulating neonatal cortical surface neurodevelopmental trajectories, conditioned on continuous age phenotypes: postmenstrual age at scan (PMA) and gestational age at birth (GA). To achieve this, we take inspiration from existing work [25], [26] and directly condition our generator on the desired age difference to learn a cortical surface difference map. Our work incorporates the following specific contributions:

- We describe a model that integrates graph convolutions into a modified CycleGAN framework that can produce fully controllable predictions across the entire neurodevelopmental trajectory of the neonates. Model design is validated through an ablation study, with hyperparameter tuning.
- We perform experiments on the developing human connectome project (dHCP) dataset [41] demonstrating the utility of our model in generating realistic, age-accurate and subject-specific predictions of cortical maturation and the effect of changes in degree of prematurity.
- We demonstrate the utility of our method by identifying biomarkers that correlate to improved predictions of cognitive outcomes of preterm subjects.

## II. Methods

### A. Problem Statement

We denote a neonatal cortical surface feature map as ***x***_*s,b*_, for a subject with PMA = *s*, and GA = *b*. We define cortical neurodevelopment as an additive process, and model changes in *s* and *b* separately:

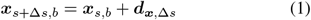

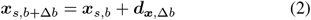

where, ***d***_***x***,Δ*s*_ is a difference map on ***x*** due to a Δ*s* change in *s*. We use **bold** notation to denote vectors throughout. The aim of our model is to learn ***d*** from ***x***, by conditioning the generator and discriminator on age values (*s,b*) and age difference values (Δ*s*, Δ*b*) as detailed in section II-D. Learning age difference maps, over directly aging images, was found to be advantageous as it helps to preserve subject identity, since intrasubject differences, due to age, are expected to be much smaller than intersubject variation due to different cortical folding patterns. This was verified this through ablation (Sec IV-A).

### B. Graph Convolutional Networks (GCNs)

The traditional form of convolution used in 2D and 3D imaging is invalid on cortical surface meshes due to their irregular local structure and the need for rotational, not translational, equivariance to correctly represent positional variation of features. In this paper we propose to use a form of graph convolutional operator which is invariant to the numbering and ordering of vertices [42]:

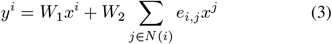

Here, *e*_*i,j*_ is the adjacency matrix, *W*_1_, *W*_2_ are learned filter weights, *x*^*i*^ denotes the *i*th vertex of ***x***, and ***y*** is the output of the convolutional operation. Since the sum operation is applied over all the neighbours of point *i* before a single learned weight (*W*_2_) is applied, the output of the network is unchanged by variation in either the number or ordering of the neighbours.

### C. Proposed Model

Continuous modelling of neurodevelopmental phenotypes is implemented through the training of an *n*-step cycle GAN. At each step (Fig 1A) two operations are implemented: 1) an age transformation operation (blue arrow) that uses the generator to age the input image by a specified age difference; and 2) an image discrimination operation (red arrow) that evaluates the accuracy of the simulated ageing through a binary cross entropy loss (*L*_*AGE*_) estimating the probability that the synthetic image is a realistic image of the target age. The network uses a closed cycle of *n* of these operations, where the image is transformed to a number of intermediate ages before closing the cycle, by transforming the image back to the original age and evaluating a reconstruction loss *L*_*RECON*_ that enforces cycle consistency i.e that the final image ***x***_*n*_ is equivalent to the initial image ***x***_0_. This takes the form of a smooth L1 loss:

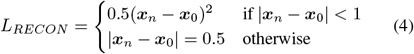

Examples of 2, 3 and *n*-cycle cases are shown in Fig 1B.

**Fig. 1:**
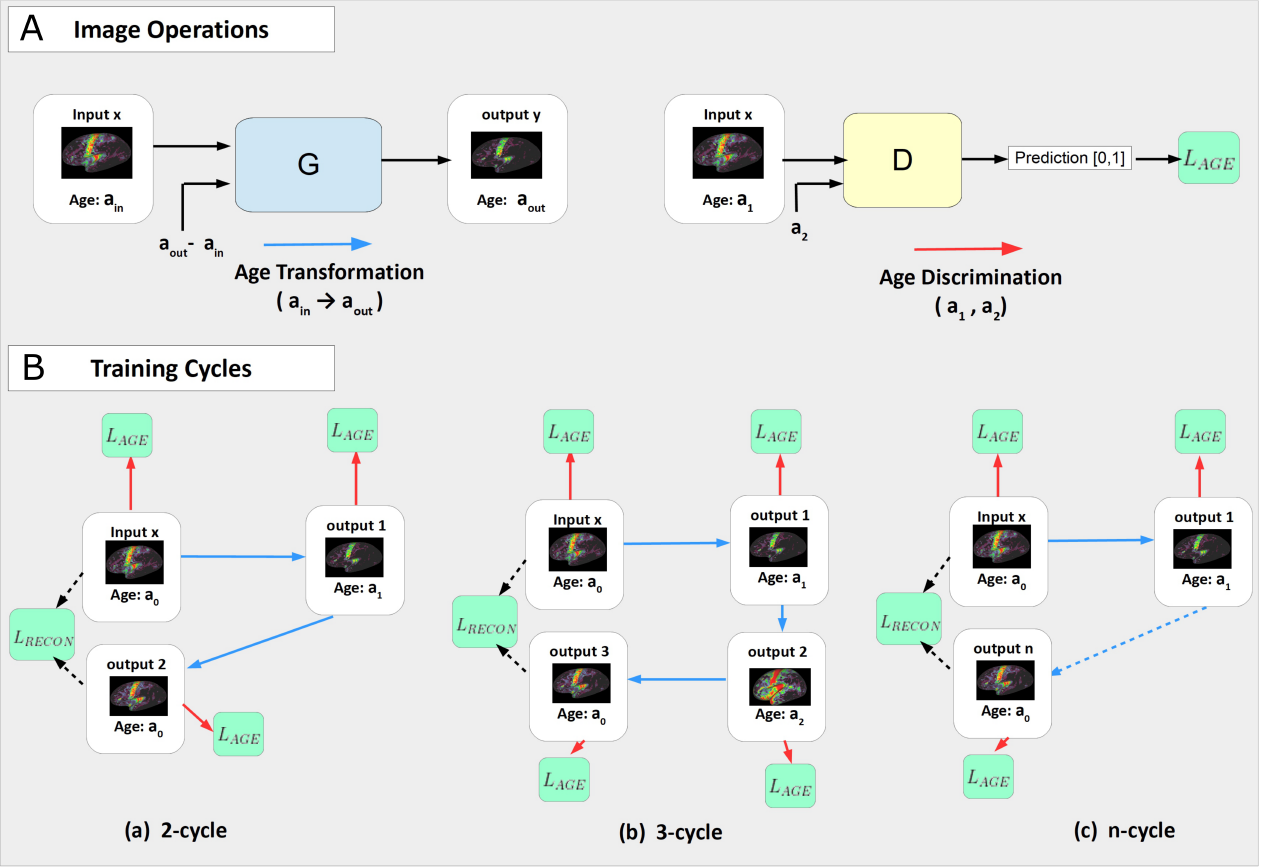
The proposed model consists of a *n*-step closed training cycle (B) of image operations (A) regularised by a cycle loss. During each step in a closed cycle, an input image is transformed to a new age (blue arrow) and an age loss is evaluated (red arrow) to measure the accuracy of the new image to the target age. The number of transformations, *n*, in a cycle is a hyperparameter. Training cycles of lengths 2, 3 and *n* are shown.

### D. Architecture

The architecture of the model is shown in Fig 2. In each case, the conditional generator and discriminator are graph convolutional networks (GCNs), with convolutions implemented using Eq 3 and convolutional layers interleaved with ReLu activation functions. The input data are sphericalised icosahedral cortical feature maps of resolution 40962 vertices (see section III-A). These are downsampled between convolutional layers, to lower resolution icospheres of resolution 10242, 2562 and 642 vertices, using an efficient hexagonal pooling scheme [36]. Upsampling of the decoder network is implemented using bilinear interpolation.

**Fig. 2:**
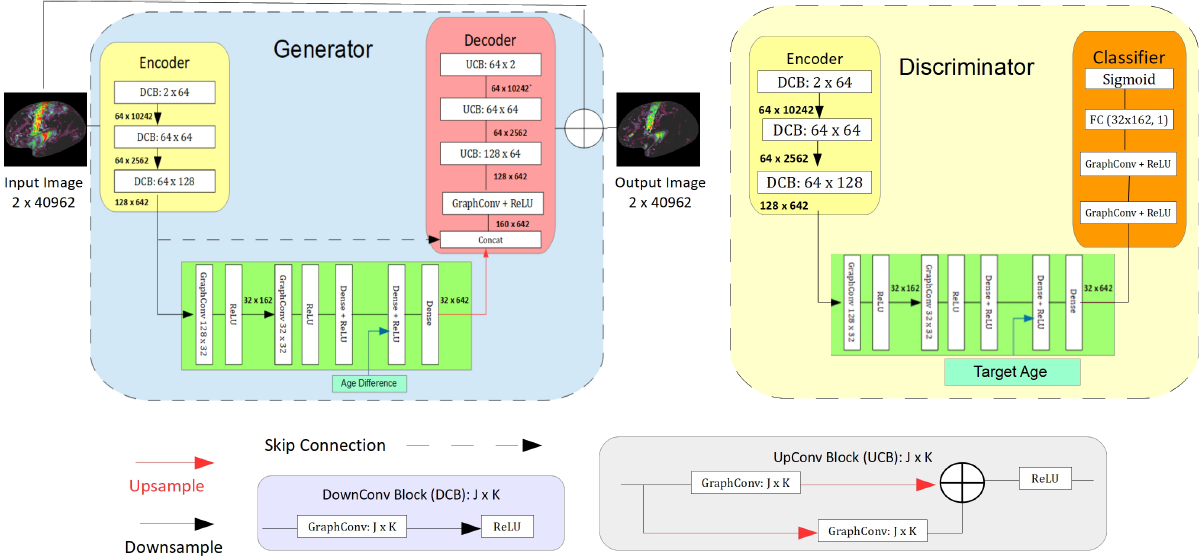
The generator consists of three sections: an encoder (yellow), an age modifier (green) and a decoder (red). The encoder performs feature extraction on an input image. The age modifier takes in the desired age difference and transforms the latent vector to one of the desired age. Original and transformed feature vectors are then combined by concatenation and passed to the decoder, which generates a difference map, adding this to the original image to obtain the transformed image. The aim of the discriminator is to determine if the input image is a realistic image of a specific target age. It takes the form of a GCN conditioned on a specific target age, that generates a probability that the input image corresponds to a real image of the target age.

The generator is composed of an encoder, age modifier component, and a decoder. The encoder extracts latent features, which the age modifier transforms according to the conditioned age difference (Δ*b* or Δ*s*), and the decoder generates an age difference map.

The discriminator takes the form of a GCN classifier, which is conditioned on a target age (*s*, or *b*) as shown in Fig 2. The output is a probability representing the confidence that the input image is a realistic image of that specific target age. In both the generator and discriminator, the conditioned variables, age difference and target age, are represented as floats, in contrast to the discrete ordinal encodings approach of prior methods [25], [26]. This allows our model to perform age progression or age regression depending on the sign of the age difference, and is intuitive as neural networks are known to (mostly) approximate smooth continuous functions [43], where we expect the modification of the image encoding to smoothly vary with age.

## III. Experimental Setup

### A. Data and Augmentations

#### 1) Dataset

All experiments were run using neuroimaging data from the publicly available dHCP dataset^*^ [1], [44]– [48]. The dataset comprises preterm and term subjects, cross-sectionally scanned between 24-45 weeks post menstrual age (PMA), and gestational ages at birth (GA) between 21-40 weeks. Table I describes the dataset and the subsets used in this paper. In the dHCP, the preterm group typically undergo a scan either shortly after birth, or at term-equivalent age, but a subset of 45 subjects are scanned twice for which longitudinal data is available.

**TABLE I:** Breakdown of the datasets used in this paper. GA and PMA measured in weeks. Figures show mean and standard deviation.

| Dataset | Count | Preterms |  |  |  | Terms |  |  |  |  |
| --- | --- | --- | --- | --- | --- | --- | --- | --- | --- | --- |
|  |  | PMA | GA | QChat | Bayley | Count | PMA | GA | QChat | Bayley |
| Full | 161 | 37.0 ± 4.0 | 31.9 ± 3.5 | 31.0 ± 10.2 | 99.7 ± 12.2 | 419 | 41.1 ± 1.7 | 40.0 ± 1.2 | 30.4 ± 8.9 | 100.4 ± 10.9 |
| Exp1 | 111 | 33.96 ± 2.27 | 31.7 ± 3.37 | 32.0 ± 9.7 | 99.4 ± 11.9 | " | " | " | " | " |
| Exp2 | 95 | 40.75 ± 2.08 | 32.06 ± 3.65 | 29.8 ± 10.7 | 100.1 ± 12.6 | " | " | " | " | " |
| Longitudinal | 45 | 33.8 ± 2.0 (first scan)<br>41.2 ± 1.4 (second scan) | 31.0 ± 3.3 | 30.7 ± 9.3 | 99.7 ± 10.8 | N/A | N/A | N/A | N/A | N/A |

Two different cognitive outcome measures were available for each subject, measured at around 18 months (592 *±*74 days at assessment): QChat [49] and Bayley-III [50]. QChat is a self-reported questionnaire based metric that assesses Autism Spectrum Disorder risk, whereas Bayley-III is a composite metric of language and motor development, derived from behavioural observations. These are also shown in table I.

#### 2) Preprocessing

The full details of image reconstruction and preprocessing pipelines are described in [1], [47] and references therein. In brief, motion corrected and reconstructed T2w and T1w sMRI images were passed through the dHCP structural pipeline^†^ [47], which performed Draw-EM tissue segmentation [51], surface extraction [48] and inflation [27] to return vertex-matched inner (white matter), outer (pial), midthickness, inflated and spherical surfaces. This process generated a number of univariate surface feature maps of which we use sulcal depth and cortical myelination defined as T1w/T2w ratio maps [52]. Individual sphericalised maps were then registered to the dHCP 40-week neonatal symmetric cortical surface template [53] using multimodal surface matching (MSM) [54], [55], driven by sulcal depth features. Aligned features were then resampled to a regular 40, 962-vertex icosphere using barycentric interpolation, implemented using Human Connectome Project (HCP) workbench software [56].

### B. Implementation

The data were augmented offline with non-linear warps to produce realistic variations in the data and improve network generalisation. These were produced by the scheme described in Fawaz, Williams, Alansary, *et al*. [36]. We used the myelination and sulcal depth features maps as our input channels. These were normalised to a mean of 0 and a standard deviation of 1. Train, validation and test sets were split evenly between preterms and terms, with ratios 0.8:0.1:0.1. Target ages for simulation were sampled at random from between 28 and 45 weeks. Graph convolutions were implemented using PyTorch Geometric [57]. Models were trained with an Adam Optimiser with learning rate of 0.001 for 100 epochs. Our code can be found at https://github.com/Abdulah-Fawaz/continuous_cGAN.

### C. Experiments

In this paper, we evaluate our proposed model through two experiments. The first to simulate changes with respect to healthy cortical maturation during the third trimester, and the second reflects the impact of prematurity on cortical neurodevelopment.

#### 1) Simulating Healthy Cortical Neurodevelopment

In our first experiment we make the simplifying assumption that scans acquired from preterm neonates, shortly after birth, are approximately healthy. We then model the effect of increasing PMA by training on cross-sectional data acquired from all preterm neonates’ first scans, and all scans from term controls. For this dataset (described in table I), it is assumed that PMA *≈* GA; thus, GA need not be modelled explicitly, and Δ*b* can be set to zero in Eq (1). To train the model on this dataset, the generator and discriminator are conditioned on Δ*s* and *s* respectively.

#### 2) Simulating the Impact of Preterm Birth

The aim of the second experiment was to simulate the impact of GA, and therefore degree of prematurity, from scans acquired around term-equivalent age. Preterm neonates first scans were excluded from the data, resulting in the dataset shown in table I. While changes with respect to healthy cortical maturation are known to be relatively stark and well-characterised in the literature [3], [6], the effect of preterm birth is relatively subtle and heterogeneous [17]. Further, the effect observed in the dHCP data set is confounded by individual’s PMA at scan. To account for both varying GA and PMA, the experiment is therefore modified to add PMA to both the generator and discriminator as a confound (concatenated as a float). Training is then conditioned on *b*, and Δ*b*. Additionally, a final information preservation loss *L*_*IP*_ is applied between successive images to encourage the network to learn changes that are proportional to the difference in GA:

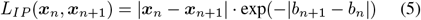

where image ***x***_*i*_ has GA of *b*_*i*_. This is to reflect that the effect of preterm birth on the brain (varying GA at birth whilst holding PMA constant) is smaller than the variation due to changes in PMA at scan for constant GA.

### D. Evaluation

For quantitative evaluation, three different metrics were used:

a. Age MAE’ - or the mean absolute error (MAE) between the apparent age of a synthetic image and the targeted age. Here apparent age was determined from an independently trained gDL regression model, implemented with 4 layers of MoNet [58] convolutions, ReLU activations and a final fully connected layer (previously described in [36]). Training was performed on the same train/val/test splits as for the proposed image generation model. This returned a baseline age MAE (for ground truth data) of 0.68 *±* 0.19 for PMA and 1.5 *±* 0.38 weeks on the more difficult task of predicting GA.
b. ‘Subject specificity’ - this evaluates the similarity between an original input scan and synthetic (aged) images. This was measured from three similarity metrics: the peak signal-to-noise ratio (PSNR), the mean square error (MSE), and the structural similarity index measure (SSIM). Note that SSIM was modified to function on the icosahedron by adapting the requisite Gaussian filtering operations to the surface.
c. Prediction of preterm outcome - clinical utility of the model of healthy cortical ageing was validated for prediction of cognitive outcome at 18 months. Here, a metric summarising deviation from typical development was derived by estimating the mean square error (MSE) between each individual’s ground truth follow up scan, and a simulated ‘healthy’ scan at the same age. This was compared against QChat and Bayley-III test scores. Since GA alone is already correlated with cognitive outcomes for preterm subjects, we measure the changes in this correlation from a GA-only baseline, to linear regression of these scores, estimated from a combination of GA + deviation metric.

### E. Ablation

Prior to running all experiments we ran ablation analyses, on the task of simulating healthy cortical neurodevelopment. Experimental design was validated against a baseline CycleGAN, trained to translate examples between two discrete classes: term (PMA *≥* 37 weeks) and preterm (PMA *<* 37 weeks) [40]. Model parameters were then evaluated by investigating the impact of changing training cycle length (from *n* = [2, 3, 4, 5]), and evaluating how well the model performed when trained to learn images directly instead of age difference maps - to test the hypothesis that learning difference maps improves subject specificity (run for the n=3 cycle only).

## IV. Results

### A. Ablation Study

The results of the ablation study are shown in table II. The CycleGAN baseline produced a high subject specificity, but a very poor age accuracy (7.66 weeks MAE) which is expected due to the discrete nature of the image conditioning. The model with the highest age accuracy (0.8 weeks MAE) was the 4-cycle model, but it also demonstrated the lowest subject specificity. The *n* = 3 model gave the next best performance in age accuracy (1.0 weeks MAE) whilst retaining the overall highest subject specificity, and thus was the best performing model overall. The same model, without difference maps, witnessed a large drop in subject specificity, confirming the hypothesis that generating translations using difference maps significantly preserves individual’s cortical morphology without compromising on the precision of the age generation. Going forward all experiments therefore use the 3-cycle network and generate difference maps.

**TABLE II:** A table comparing the performance of different generative models on the simulation of typical cortical developmental trajectories, as measured by the accuracy of synthesised images to target PMA and subject specificity.

| Model | Age Accuracy<br>(MAE / weeks) | Subject Specificity |  |  |
| --- | --- | --- | --- | --- |
|  |  | SSIM | PSNR | MSE |
| CycleGAN [40] | $7.66 \pm 2.11$ | 0.80 | **25.6** | 177 |
| 2-cycle | $2.70 \pm 0.70$ | 0.74 | 17.3 | 203 |
| 3-cycle | $1.02 \pm 0.2$ | **0.82** | 17.9 | **167** |
| 3-cycle w/o diff maps | $1.14 \pm 0.46$ | 0.66 | 13.2 | 251 |
| 4-cycle | **$0.8 \pm 0.38$** | 0.70 | 16.1 | 231 |
| 5-cycle | $1.55 \pm 0.50$ | 0.74 | 16.1 | 208 |

### B. Simulating Healthy Cortical Development

On simulation of healthy cortical neurodevelopment, data generated from the network returned an age MAE of 1.02 *±* 0.2 weeks. This compares well against the performance of the baseline regression network on ground truth data (Fig. 4) with some greater error at the extremes of the distribution, where fewer data samples were available.

**Fig. 3:**
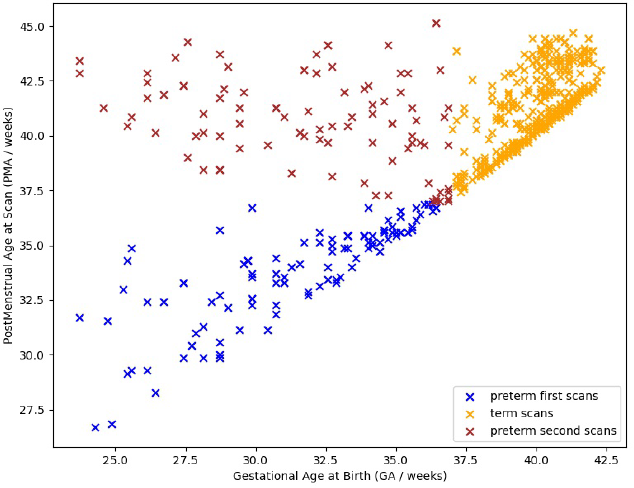
A plot to show the distribution of images in our dataset.

**Fig. 4:**
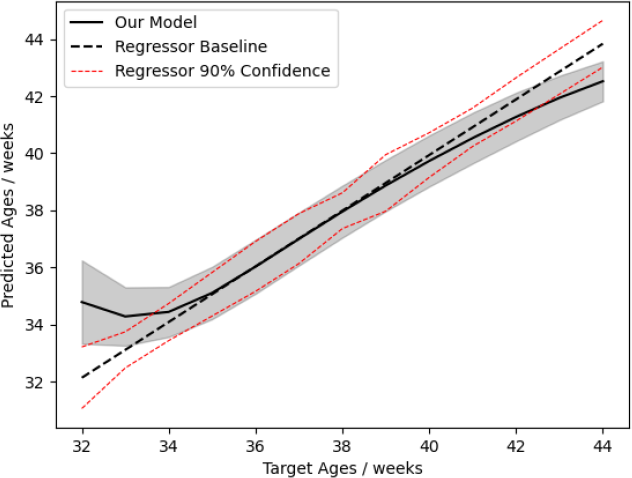
A comparison of age MAE for simulated and ground truth cortical feature maps. Here the X-axis reports target (or ground truth) PMA and the Y-axis reports PMA predicted from the baseline regressor. Performance on generated images is shown with a solid line, compared to ground truth (dashed). The grey region indicates the uncertainty in the age accuracy of the GCN model on generated images, and the red dashed lines show the 90% confidence interval of the regressor.

Examples of the images generated by the model across time are shown in Fig. 5 for two example preterm individuals. It can be seen, through comparisons to population average templates [53], [59], that changes in myelination (top) and sulcal depth (bottom) follow expected neurodevelopmental trends. Myelination increases most strongly along the central sulcus and major sulci deepen and grow as PMA increases. More subtle features, such as myelination of the middle temporal (MT) area, and the emergence and growth of smaller sulci along the temporal lobe are present. However, this cannot simply be explained as the network generating a population average model of development. Visual inspection of the cortical folding patterns for each subject show highly divergent patterns of folding in the frontal lobe which are preserved during simulated ageing. Follow up scans are shown for reference, but these are not expected to match precisely as our model is only simulating healthy cortical neurodevelopment without taking into account the impact of preterm birth. This property of identity preservation was further investigated through statistical comparison of each preterm individual’s follow up scan with their simulated images. The mean square error (MSE) difference between the sulcal depth feature maps of a subject’s follow up scan and its own simulated image was lower than any other subject (aged to the same PMA) for 42 of the 45 subjects for which longitudinal data was available (93.3%). The average margin between the correct subject’s generated image and the next-closest subject was 18.4 *±* 13.3 MSE. Sulcal depth features maps were used because they contain an individual’s unique folding pattern, and are therefore a good marker of subject preservation.

**Fig. 5:**
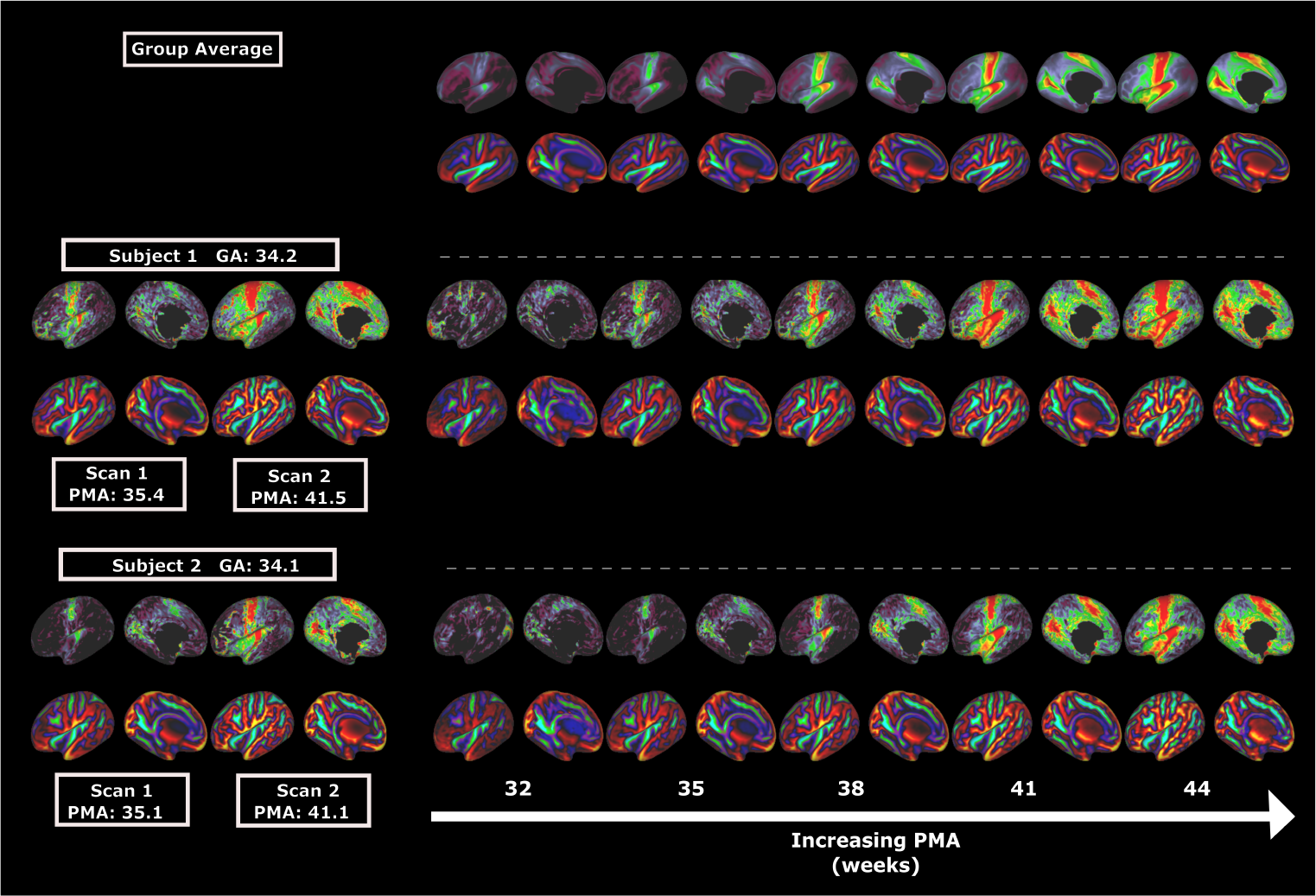
Group average templates for the dHCP dataset (top), a subject of GA 34.2 weeks, and scans at PMA 35.4 weeks and 41.5 weeks (middle, left) and a second subject of GA 34.1 weeks and PMA 35.1 weeks and 41.1 weeks (bottom, left) alongside a set of sample synthetic images of (columns 2-6) generated by the model to represent the same subjects at PMA 32, 35, 38, 41 and 44 weeks respectively. The model predicts an increase in myelination and a change in sulci depth consistent with those seen in the group average templates, while preserving individual cortical folding patterns.

Further, we find that the proposed biomarker (MSE deviation between ground truth and simulated follow up scans) increases the *r*^2^ correlation between predicted and ground truth 18-month cognitive test scores (for the same 45 left out individuals) with QChat correlation increasing slightly from 0.30 to 0.32, and the Bayley-III correlation increasing more substantially from 0.41 to 0.53. Poorer performance on the Qchat may be explained by the fact that it is primarily used for screening ASD, but there are no-known individuals at high risk of ASD in the dHCP dataset [1].

### C. Simulating the Impact of Preterm Birth

Age MAE for the second experiment was 1.3 *±* 0.8 weeks. While this is larger than for the PMA experiment, it remains close to the 1.5 *±* 0.38 obtained from ground truth data (Fig 7) indicating that the performance of the regressor is likely a limiting factor in evaluating the true performance of the generative model. Nevertheless, there is a clear trend that points to our model being able to monotonically alter the apparent GA with target GA.

Examples of the images generated by the model for different GA are shown in Fig 6. The first row shows an example term-control subject with GA 40.8 weeks (PMA at scan 41.0 weeks) as the model simulates decreasing GA to 32 weeks. Conversely, the second example shows a preterm subject with GA 28.7 weeks and PMA at scan of 43.7 weeks, altered so that their GA appears 40 weeks.

**Fig. 6:**
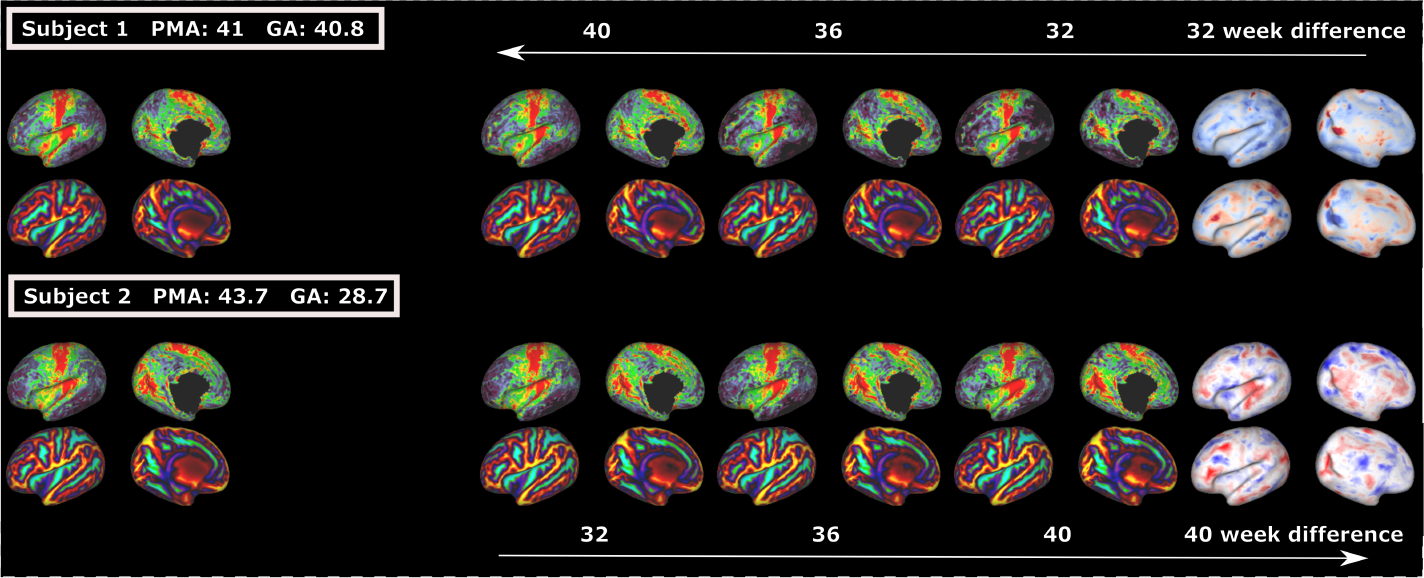
(Top) A subject of original GA 40.8 weeks and PMA 41.0 weeks with simulated decrease of GA to 40, 36 and 32 weeks. (Bottom) A preterm subject of original GA 28.7 weeks and PMA 43.7 weeks aged up to a GA of 40 weeks. Images in the final column show difference maps between the original images and the final synthetic images.

**Fig. 7:**
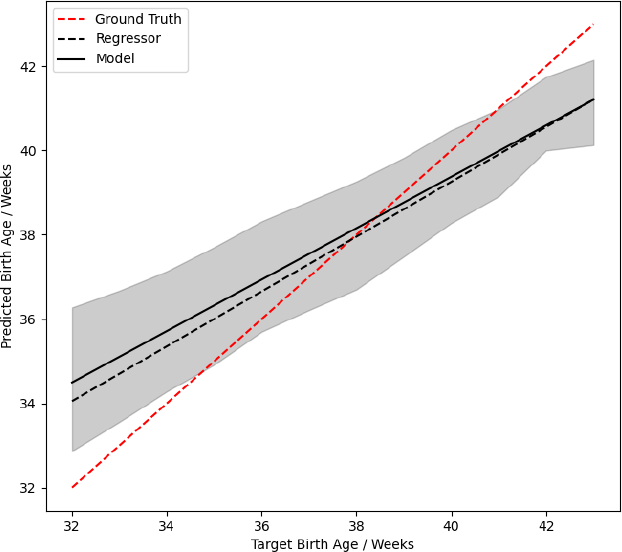
A plot showing the predicted gestational age (GA) vs input target age of our GCN model (solid) compared against the baseline regressor accuracy (dashed). The grey region indicates the standard error of the GCN model predictions, and the red dashed line shows the ground truth.

In contrast to the rapid changes that occur as PMA is increased from 32 to 44 weeks, differences in GA are associated with more subtle changes. To aid interpretation, the final column shows difference maps between the original images and the final synthetic images. These show a global reduction in myelination across most of the cortical surface as GA decreases - in line with a large body of existing research that links preterm birth with delayed myelination [47], [60], [61]. The decrease in myelination is not total, however, and certain regions of the preterm brain are known to display higher levels of myelination suspected to be due to early exposure to stimuli [62], and these too are present in our difference maps.

The impact of decreasing GA on sulcal depth is less well-documented, but is known to impact shallowing of the sulci and gyri [6] and this is reflected in the correspondence between the folding pattern and the difference maps. Regional analysis [3], [6] of the cortical surface of preterms identified relatively shallow temporal and cingulate sulci as regions most consistently affected by preterm birth - the two regions most strongly highlighted by our model.

## V. Conclusion

Deep generative modelling on surfaces presents challenges on two fronts. Modelling of surfaces is complicated by the lack of a global coordinate system over the surface on which to define convolutions [63], and difficulty defining regular up/down sampling [29]. Deep generative models must contend with the limited availability of paired data in medical imaging, retaining subject identity during image generation, whilst simultaneously modelling cortical heterogeneity across populations.

In this paper, we have presented a novel deep generative model of neonatal cortical surface neurodevelopment that solves both sets of challenges by integrating state of the art methods from geometric deep learning and Euclidean generative modelling. The model was thoroughly validated through quantitative and qualitative assessments, focusing on the accuracy of age prediction for generated images, and the preservation of subject identity. Our results demonstrate the success of our model in predicting changes associated with cortical maturation and degree of prematurity. Furthermore, the model was used to propose a novel biomarker that improved the accuracy with which it is possible to predict cognitive outcomes of at risk individuals. Longterm, such an approach could be utilised in a clinical environment to identify subjects for clinical intervention.

Moving forward, there are several opportunities for further exploration and improvement. Our model could be applied to other forms of MRI surface data beyond sMRI, e.g. diffusion-weighted imaging or functional MRI. Additionally, conducting studies incorporating larger datasets would allow us to validate our findings across diverse populations and improve the generalisability of our model, particularly as it was observed that our model accuracy decreased in regions of lower data availability.

More fundamentally, there was an underlying assumption that scans acquired from preterm births near the time of birth were healthy. This is highly unlikely to be satisfied in practice since underlying risk factors, genetic and environmental, are likely to be implicated in the occurrence of preterm birth. In future, this assumption could be relaxed with the use of fetal scans to represent healthy subjects at lower PMA. Another avenue of further work, inspired by Xia, Chartsias, Wang, *et al*. [26], would be to condition the model on additional discrete factors such as disease state, sex, or birth weight, to further disentangle how demographic and clinical factors impact neurodevelopment.

## Footnotes

* http://www.developingconnectome.org

† https://github.com/BioMedIA/dhcp-structural-pipeline

